# Multimodal functional characterization of the *LDLR* noncoding landscape

**DOI:** 10.64898/2026.10.22.761041

**Authors:** Hitarthi S. Vyas, Kyle Leix, Bala B. Burugula, Sanjana M. Paye, Chinmay Raut, Elizabeth K. Speliotes, Brian T. Emmer

## Abstract

The expression level of the low-density lipoprotein receptor (*LDLR*) gene is a critical determinant of circulating low-density lipoprotein concentrations and the corresponding risk of atherosclerotic cardiovascular disease. While the importance of sterol response elements in *LDLR* transcriptional regulation has been firmly established, the functional contribution of other potential *cis*-regulatory elements (CREs) remains poorly understood. In this study, we developed and applied a variety of high throughput functional genomic tools to systematically interrogate the noncoding space at the *LDLR* locus. These included saturation CRISPR nuclease, CRISPR activation, and CRISPR inhibition screening as well as a massively parallel reporter assay. With each approach, we observed robust discrimination of control sequences and significant effects for other regions with a previously unrecognized influence on *LDLR* expression. The CREs identified by each modality were overlapping but distinct, both in relation to each other and to those regions predicted to harbor CRE activity. Collectively, our findings clarify the functional landscape of the *LDLR* locus and highlight the complementary nature of different functional genomic approaches to CRE identification.

## INTRODUCTION

The low-density lipoprotein receptor directly binds to circulating low-density lipoprotein (LDL) particles and mediates their clearance from the bloodstream, thereby reducing an individual’s risk of atherosclerotic cardiovascular disease (ASCVD). Multiple classes of therapeutics that increase hepatic LDLR activity are effective in preventing and treating ASCVD^1^ while human genetic variants associated with increased LDLR activity are associated with reduced ASCVD risk^2-4^. Despite the clinical importance of *LDLR* expression, the *cis*-regulatory elements that modulate it remain largely unmapped aside from sterol response elements in the promoter region.

High-throughput methods for CRE identification have typically involved the detection of associated features like accessible chromatin, specific histone modifications, or transcription factor binding. These approaches have nominated an estimated ∼2 million candidate CREs across the human genome, including many at the *LDLR* locus^5^. Several studies, however, have found CRE prediction based on biochemical or sequencing features alone to be imprecise, underscoring the importance of functional testing^6^. Historically, empirical testing has typically been restricted to one or a few candidates at a time, but advances in genome engineering and massively parallel sequencing have greatly increased the scalability of functional assays. High-throughput approaches now include pooled CRISPR nuclease (CRISPRn) screening, in which indel formation at candidate CREs may disrupt the recruitment of *trans*-regulators of gene expression, as well as CRISPR inhibition (CRISPRi) and activation (CRISPRa) screening, in which candidate CREs may be perturbed by the recruitment of CRISPR-associated enzymes linked to modulators of transcriptional activity. Similarly, in a massively parallel reporter assay (MPRA), the activity of candidate CREs may be assessed by measuring how they affect the expression of barcode-associated reporter transcripts. While several studies have successfully applied these approaches to interrogate CREs of different genes, to our knowledge none have applied all of them to the same noncoding space.

We previously performed CRISPR nuclease screening for candidate *LDLR* CREs^7^. We identified 25 regions whose disruption reduced *LDLR* expression and, for one of these, we mapped enhancer activity to a discrete 129 bp element in the first intron. Motivated by the technical performance of this screen together with uncertainty in its reproducibility and convergence with other approaches for CRE identification, in this study we deepen our investigation of *LDLR* CREs with saturating testing by parallel CRISPRn, CRISPRi, and CRISPRa screens and MPRA. We identify several previously unrecognized *LDLR* functional elements and provide insights into the overlapping but distinct information provided by each approach.

## RESULTS

### Expanded CRISPR nuclease screening for *LDLR* CREs demonstrates reproducibility and identifies new functionally important regions

To first validate our prior CRISPR nuclease screen of candidate *LDLR* CREs and to expand our interrogation to a broader genomic space, we designed and synthesized a custom library of 16,624 gRNAs targeting every SpCas9 PAM motif in a 124 kb region extending from 50 kb upstream of the *LDLR* transcription start site (TSS) to 30 kb downstream of its 3’ UTR (Fig 1, Supplementary Table 1). We then delivered each gRNA in a pooled fashion at a low multiplicity of infection (MOI) to an average of >500 cells apiece expressing a wild-type SpCas9 construct that conferred robust disruption of LDL uptake when combined with a control *LDLR* CDS-targeting gRNA (Fig S1A). We identified regions influencing *LDLR* expression by quantifying their relative gRNA abundance in FACS-isolated 10% subpopulations with the most and least uptake of exogenous fluorescently conjugated LDL. Consistent with our prior targeted CRISPR nuclease screen, this approach again demonstrated excellent technical performance, as reflected by clustering of samples by sorting bin on principal component analysis (Fig S2A), clear discrimination of control gRNAs (Fig 2A), and consistency of control gRNA effects across independent biologic replicates (Fig S2D). As expected, we observed the strongest activity for gRNAs targeting either *LDLR* coding sequences or noncoding regions in proximity to intron-exon junctions (Figs 2A, S2E). As in our prior nuclease screen, we therefore filtered out gRNAs whose target localized within 100 bp of an intron-exon junction to minimize confounding by indels that disrupt mRNA splicing and/or protein coding sequence.

**Figure 1.**
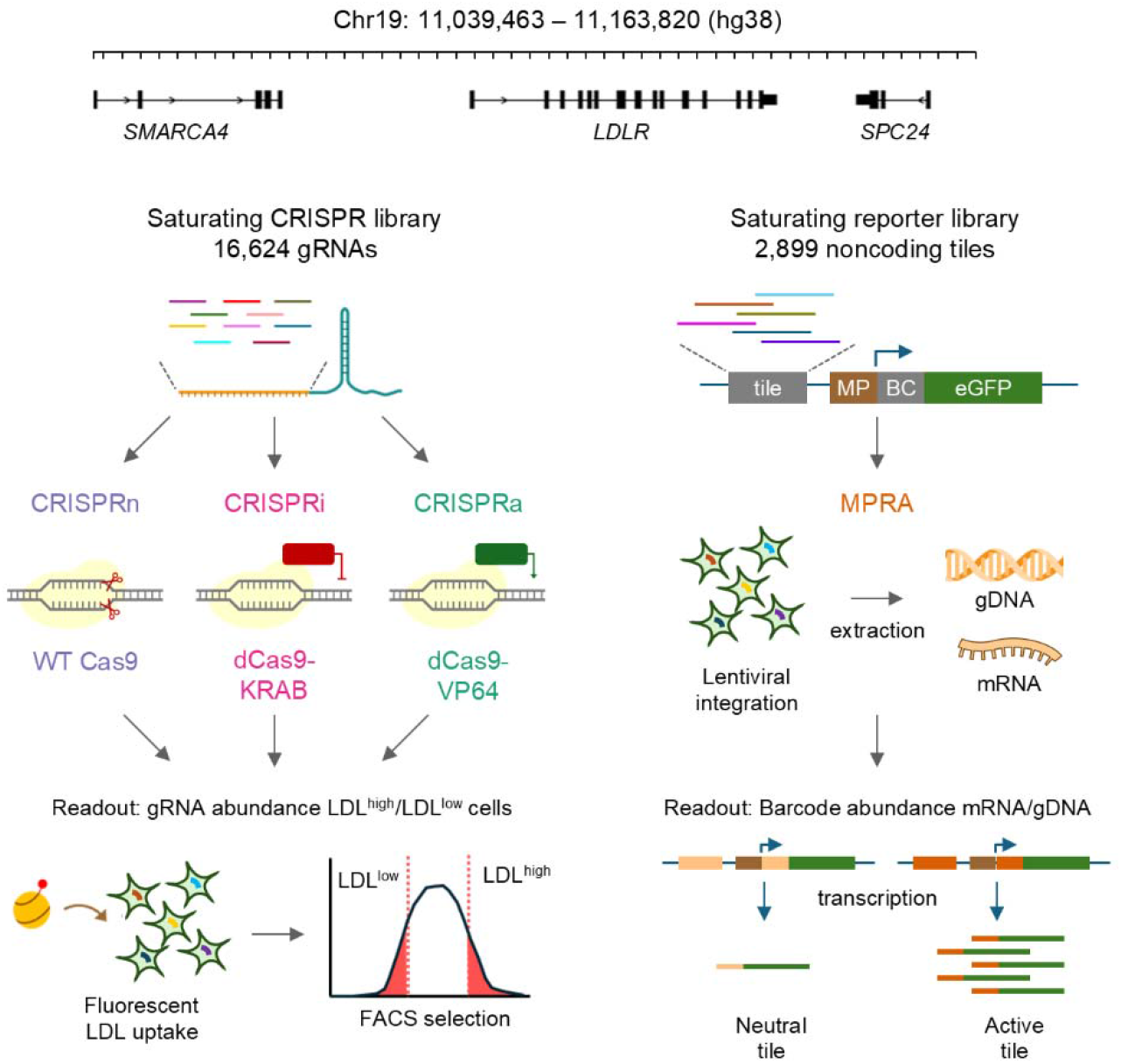
Overview of study approach. The *LDLR*-spanning region of interest was functionally interrogated by saturation CRISPR nuclease, inhibition, and activation screening and by a massively parallel reporter assay (MPRA). For CRISPR screening, a pooled gRNA library with all 16,624 SpCas9 gRNA targets was delivered to HuH7 cells in combination with Cas9 nuclease (CRISPRn) or catalytically dead Cas9 mutants fused to KRAB (CRISPRi) or VP64 (CRISPRa). The impact of each gRNA on *LDLR* expression was assessed by quantifying its relative enrichment in cells with high versus low uptake of fluorescent LDL. For MPRA screening, a pooled library containing all 2,899 overlapping 129 bp tiles plus controls was inserted along with an associated 5’ UTR barcode (BC) into a reporter construct with eGFP expression driven by a minimal promoter (MP). The transcriptional activity of each tile was assessed by quantifying the abundance of its associated BCs in mRNA relative to genomic DNA.

**Figure 2.**
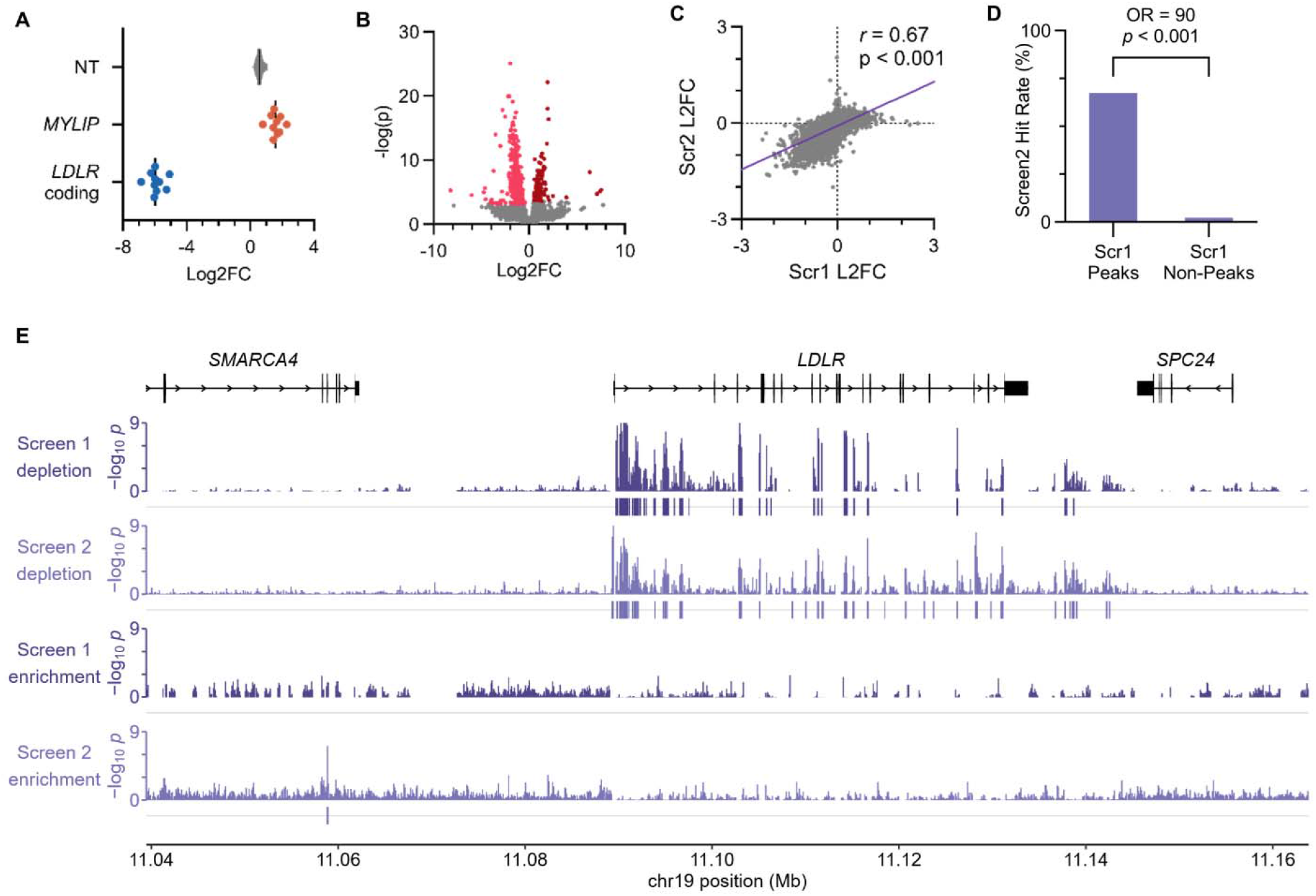
CRISPR nuclease screen. (A) Relative abundance of individual control gRNAs either lacking a genomic target site (NT) or targeting the 5’ UTR of the *LDLR* negative regulator *MYLIP* or the coding sequence of *LDLR*. Values represent the log2 fold change of gRNA counts in LDL^high^ cells relative to LDL^low^ cells. (B) Volcano plot of log2 fold change in gRNA abundance in LDL^high^ cells relative to LDL^low^ cells together with associated negative log-transformed p-values. Individual gRNAs with significant (BH-adjusted p<0.01) depletion or enrichment in LDL^high^ cells are highlighted in pink and red, respectively. (C) Correlation of LDL^high^ vs LDL^low^ log2 fold change for individual gRNAs common to both this study and our original targeted CRISPR nuclease screen of candidate *LDLR* CREs. (D) Proportion of genomic coordinates identified as significant positive regulators of LDL uptake by sliding 100 bp window analysis in this study, stratified by whether they were detected as hits or not in our original targeted CRISPR nuclease screen. (E) Genomic tracks of empirical p-values and called peaks for depletion and enrichment for all gRNAs targeting sliding 100 bp windows over the region of interest. The same pipeline was applied for both the original target screen (Scr1) and this study’s saturating screen (Scr2).

Among the remaining test gRNAs in the library, we observed many with significant depletion in LDL^high^ cells, consistent with indel formation at their target sites leading to a reduction in *LDLR* expression (Fig 2B, Supplementary Table 2). Individual gRNAs that caused a defect in LDL uptake exhibited a high degree of spatial clustering (Fig S2B). By contrast, relatively few gRNAs were significantly enriched in LDL^high^ cells with less spatial clustering among the hits (Fig S2C), suggesting a relative lack of repressive regions in the tested interval. Comparison of enrichment for individual gRNAs common to both our original candidate-based screen and this study revealed a high degree of reproducibility (Fig 2C, Pearson coefficient = 0.67, *p* <0.001).

To next smooth and aggregate individual gRNA effects across candidate CRE regions, we averaged gRNA log2 fold change values over 100 bp sliding windows centered on each gRNA target site. We calculated empirical p-values by comparison to simulations of the same number of randomly selected gRNAs and merged contiguous regions into single CRE peaks. In total, we identified 38 regions where nuclease targeting disrupted *LDLR* expression and only 1 where nuclease targeting increased expression. Applying the same sliding window pipeline to our prior nuclease screen, we found those regions identified as CREs in our prior study were far more likely than non-peaks to be identified in the present study (Fig 2D, odds ratio = 89, *p* < 0.0001). Taken together, these findings support the reproducibility of our CRISPR nuclease screening for *LDLR* CREs, validate the functional significance of several regions, and identify new regions likely to be involved in *LDLR* gene regulation.

### CRISPR inhibition screening recapitulates most *LDLR* CREs identified by CRISPR nuclease screening

Like CRISPR nuclease screening, CRISPR inhibition screening seeks to identify functional CREs by testing for a change in gene expression upon their disruption. Unlike CRISPR nuclease screening, CRISPR inhibition achieves this without generating DNA double strand breaks and indels of variable size that may confound the interpretation of gRNA effects. Instead, CRISPRi relies on the recruitment of an epigenetic inhibitor fused to a catalytically dead Cas9. To directly compare the performance of these approaches for *LDLR*, we therefore tested our saturating *LDLR* gRNA library in combination with a dCas9-KRAB fusion that effectively reduced *LDLR* expression when combined with a control promoter-targeting gRNA (Fig S1B).

As with CRISPR nuclease screening, quality control of our CRISPR inhibition screen confirmed sample clustering by principal components (Fig S3A) and consistent discrimination of control sequences across independent gRNAs (Fig 3A) and biologic replicates (Fig S3D). Analysis of individual gRNA enrichment revealed more gRNAs associated with significantly decreased *LDLR* expression than increased expression (Fig 3B, Supplementary Table 3), with significant spatial clustering only apparent for the former group (Fig S3B-C).

**Figure 3.**
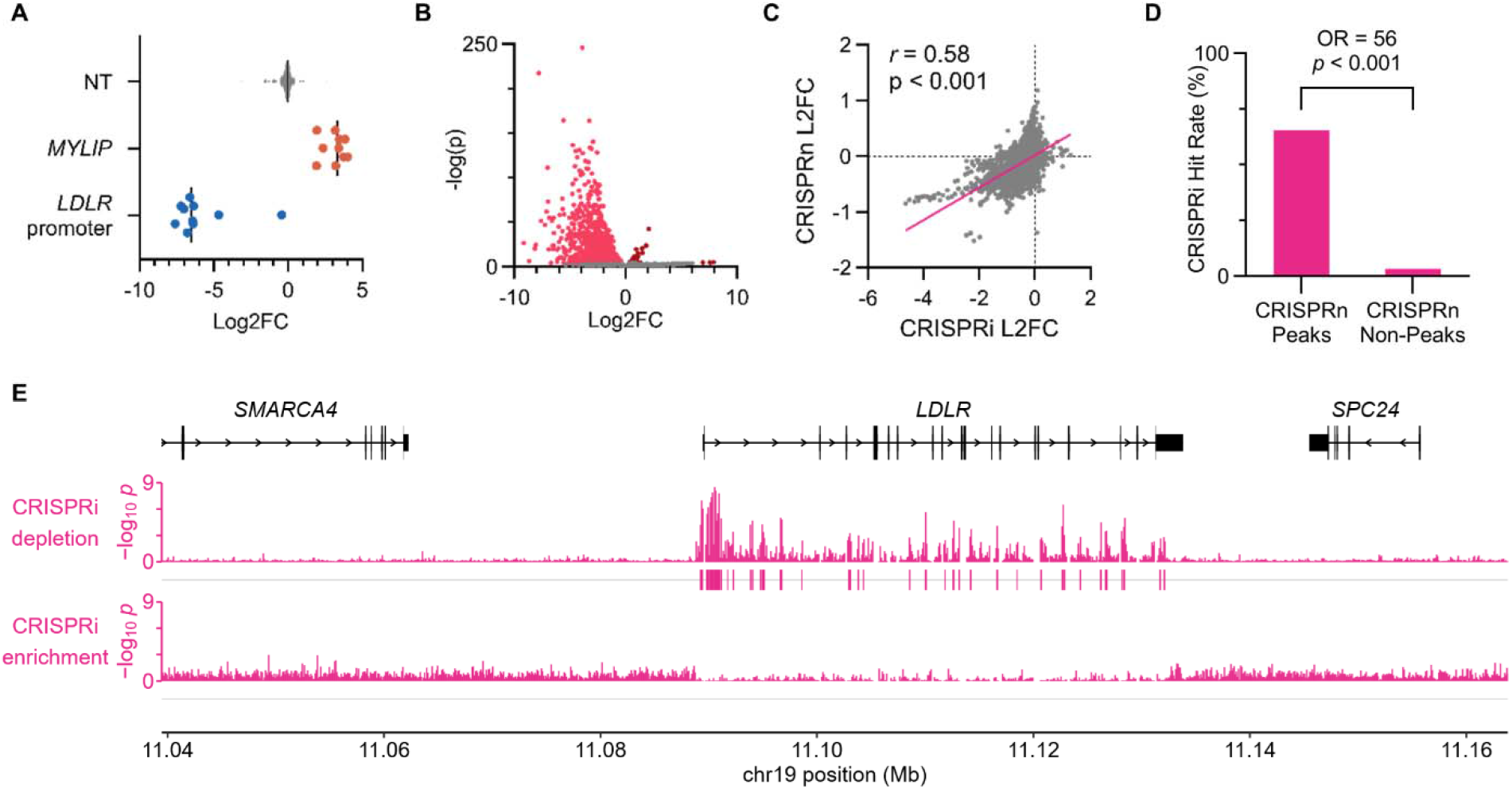
CRISPR inhibition screen. (A) Relative abundance of individual control gRNAs either lacking a genomic target site (NT) or targeting the *MYLIP* or *LDLR* promoters. Values represent the log2 fold change of gRNA counts in LDL^high^ cells relative to LDL^low^ cells. (B) Volcano plot of log2 fold change in gRNA abundance in LDL^high^ cells relative to LDL^low^ cells together with associated negative log-transformed p-values. Individual gRNAs with significant (BH-adjusted p<0.01) depletion or enrichment in LDL^high^ cells are highlighted in pink and red, respectively. (C) Correlation of individual gRNA LDL^high^ vs LDL^low^ log2 fold change in the CRISPR nuclease and inhibition screens. (D) Proportion of genomic coordinates identified as significant positive regulators of LDL uptake by sliding 100 bp window analysis in the CRISPRi screen, stratified by whether they were detected as hits or not in the saturating CRISPR nuclease screen. (E) Genomic tracks of empirical p-values and called peaks for depletion and enrichment for all gRNAs targeting sliding 100 bp windows over the region of interest.

Aggregation of gRNA effects into 100 bp sliding windows identified 34 functional CREs associated with a decrease in *LDLR* expression and none with an increase in *LDLR* expression (Fig 3E). Individual gRNA effects in the CRISPR inhibition and nuclease screens were highly concordant, with a positive correlation that was comparable to that observed between independent nuclease screens (Fig 3C, Pearson coefficient = 0.58, p< 0.001). Genomic coordinates that were identified as promoting *LDLR* expression in the nuclease screen were far more likely to be identified in the CRISPRi screen than those that were not (Fig 3D, odds ratio = 56, *p* < 0.0001). The similarity in findings between these approaches reinforces the functional significance of the identified regions and strongly suggests that the identified regions in the CRISPR nuclease screen did not represent false positives arising from larger indels that had disrupted splice sites, coding sequences, or adjacent CREs.

### CRISPR activation screening identifies *LDLR* noncoding regions that are almost entirely distinct from nuclease and inhibition screening

Whereas CRISPR nuclease and inhibition screening seek to identify CREs via their disruption, CRISPR activation tests the ability of a region to modulate gene expression when targeted for recruitment of transcriptional activators. After confirming the potential of a dCas9-VP64 fusion to increase *LDLR* expression when combined with a control promoter-targeting gRNA and a modified scaffold for additional recruitment of PP7–p65–HSF1^8^ (Fig S1C), we scaled our interrogation to the *LDLR*-saturating gRNA library.

Quality control analysis by principal components (Fig S4A), control gRNA segregation (Fig 4A dot plot), and individual replicate correlations (Fig S4D) demonstrated a technical performance for the CRISPRa screen comparable to the CRISPRn and CRISPRi screens. As expected and in opposition to the CRISPR nuclease and inhibition screens, the CRISPRa screen identified many more guides with significant enrichment rather than depletion in LDL^high^ cells (Fig 4B, Supplementary Table 4), with greater spatial clustering for the former group (Fig S4B-C). Sliding window analysis identified 19 regions where dCas9-VP64 recruitment led to a significant increase in *LDLR* expression and 3 regions with reduced *LDLR* expression (Fig 4E).

**Figure 4.**
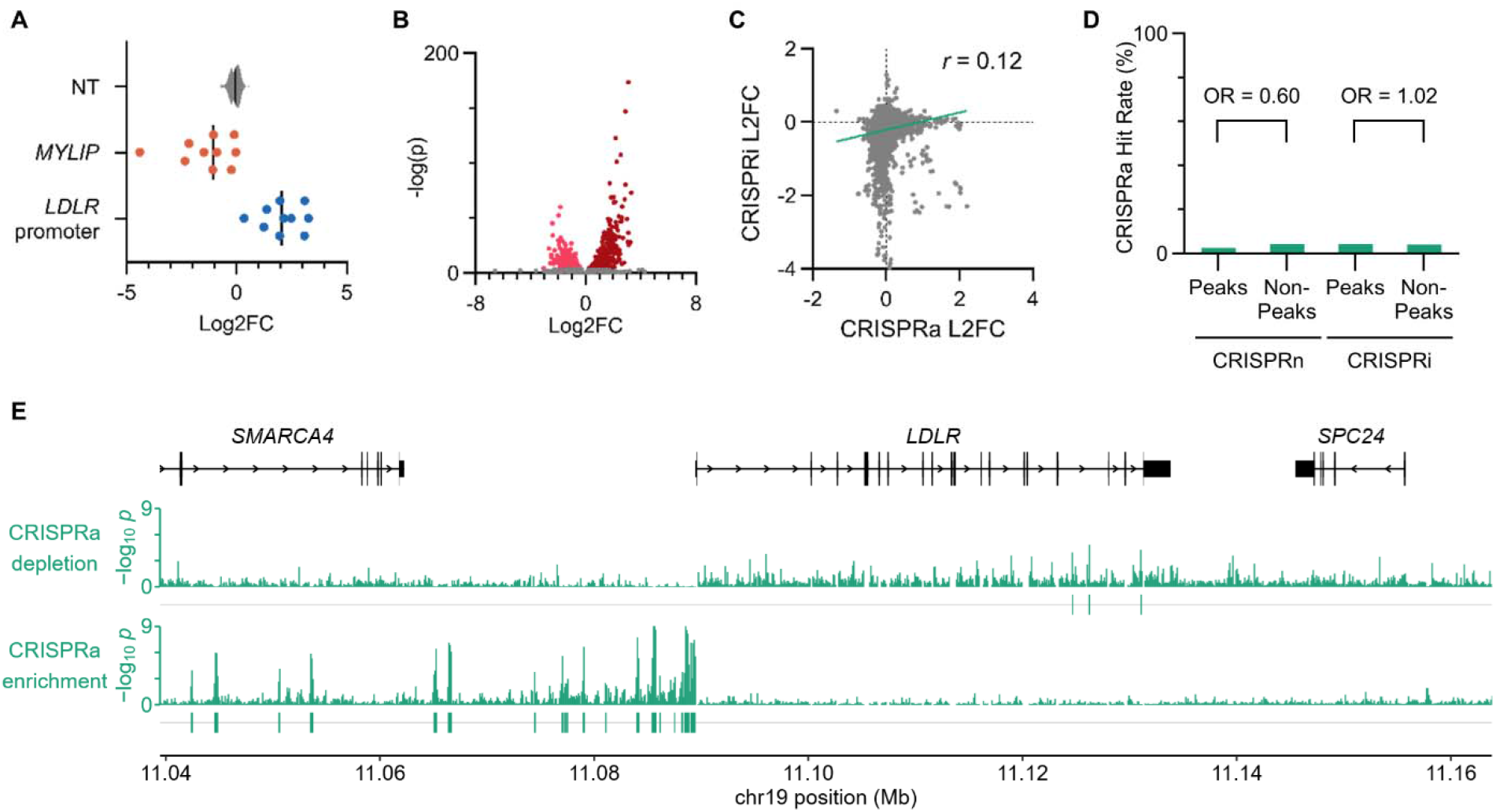
CRISPR activation screen. (A) Relative abundance of individual control gRNAs either lacking a genomic target site (NT) or targeting the *MYLIP* or *LDLR* promoters. Values represent the log2 fold change of gRNA counts in LDL^high^ cells relative to LDL^low^ cells. (B) Volcano plot of log2 fold change in gRNA abundance in LDL^high^ cells relative to LDL^low^ cells together with associated negative log-transformed p-values. Individual gRNAs with significant (BH-adjusted p<0.01) depletion or enrichment in LDL^high^ cells are highlighted in pink and red, respectively. (C) Correlation of individual gRNA LDL^high^ vs LDL^low^ log2 fold change in the CRISPR inhibition and activation screens. (D) Proportion of genomic coordinates identified as significant positive regulators of LDL uptake by sliding 100 bp window analysis in the CRISPRa screen, stratified by whether they were detected as hits or not in the saturating CRISPR nuclease and inhibition screens. (E) Genomic tracks of empirical p-values and called peaks for depletion and enrichment for all gRNAs targeting sliding 100 bp windows over the region of interest.

Visualization of CRISPRa peaks by genomic coordinate revealed a stark contrast to the CRISPRn and CRISPRi screens. While the latter approaches had identified significant regions mostly in *LDLR* introns, CRISPRa peaks were almost entirely localized in the 5’ intergenic space (Fig 4E). There was minimal correlation of individual gRNA effects by CRISPRi and CRISPRa (Fig 4C) and the identification of a genomic coordinate in a CRISPRn or CRISPRi screen peak had no significant predictive value for its identification in the CRISPRa screen (Fig 4D). The only region identified by the CRISPRa screen that was identified by either the CRISPRn or CRISPRi screens was the promoter.

### A massively parallel reporter assay identifies discrete *LDLR* sequences sufficient to drive reporter gene expression

In contrast to noncoding CRISPR screens that perturb CREs at their endogenous genomic sites, an MPRA tests the sufficiency of individual sequences to modulate the expression of barcode-tagged reporter transcripts. We designed and synthesized a library of 2,899 staggered 129 bp sequences spanning the same ∼124 kb region interrogated by the above CRISPR screens, supplemented with negative control random sequences and positive control *cis*-regulatory sequences adapted from previous MPRA studies^9^.

Analysis of the expression level associated with each tile using MPRAnalyze^10^ confirmed the reliable discrimination of control sequences, with 48/49 detected synthetic sequences that had demonstrated the most activity in a prior study similarly identified here in comparison to 0/50 that had demonstrated the least activity in the same study (Fig 5A, Supplementary Table 5). Fewer naturally occurring enhancers with previous evidence of activity in HepG2 cells showed significant activity in our study (21/50), though they were still clearly enriched relative to negative control random sequences (*p* < 0.001). The activity of all control tiles was reproducible across biological replicates (Fig S5A, Pearson coefficients = 0.81 - 0.94). Consistent with orientation-independence of enhancer activity, paired control tiles in forward and reverse orientation exhibited very similar activities (Fig S5B, Pearson coefficient = 0.91).

**Figure 5.**
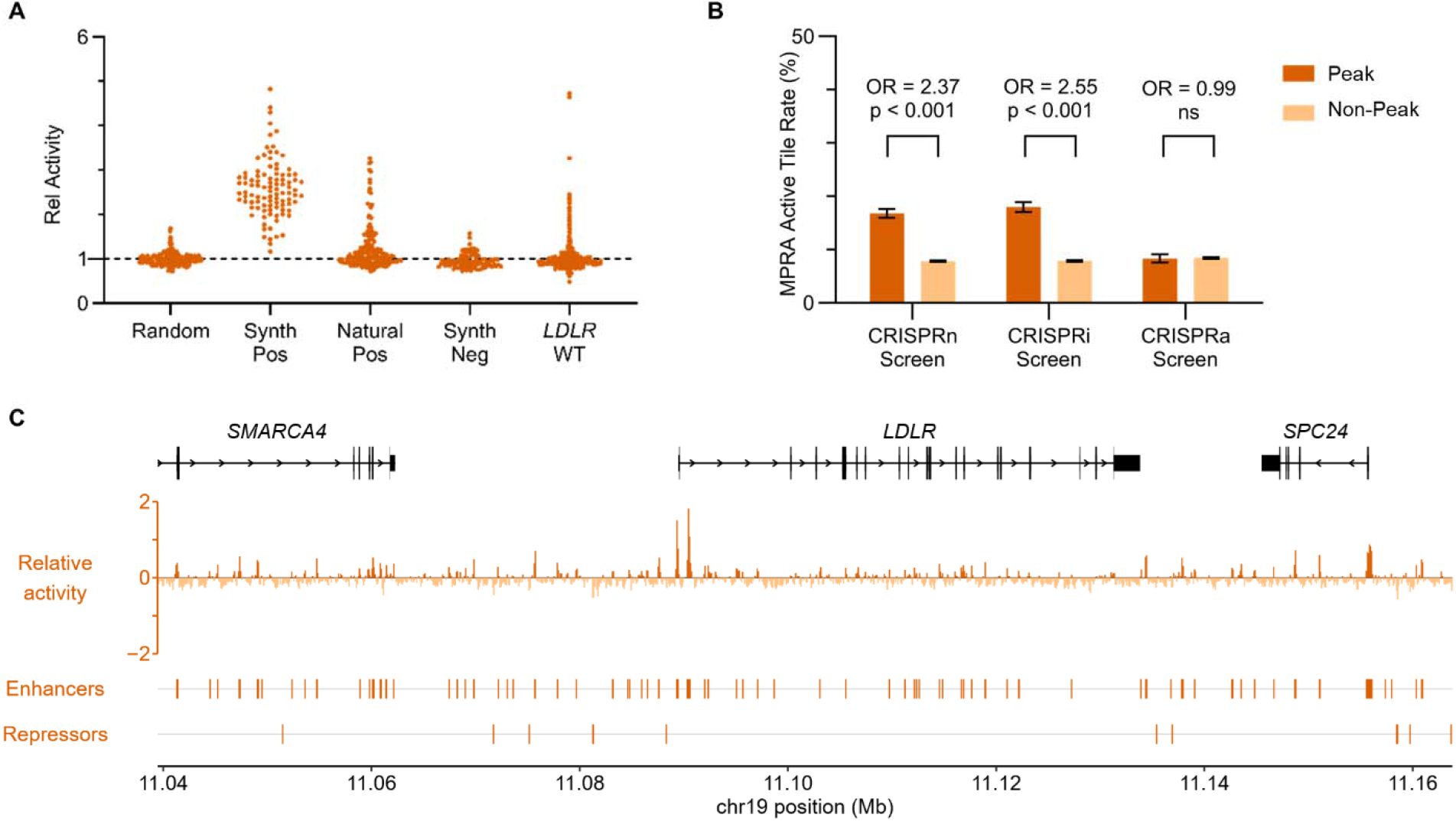
Massively parallel reporter assay. (A) Relative activity, normalized to the median of random control sequences, in HuH7 cells for each group of tiles in the library. (B) Proportion of coordinates overlapping an MPRA active tile, stratified by whether the coordinate fell within a peak of the indicated CRISPR screen or not, along with associated odds ratios. (C) Spatial distribution of MPRA activity and peak calls across the region of interest. For each coordinate, the plotted relative activity represents the average of each staggered tile overlapping it.

Among test tiles spanning the *LDLR* noncoding space, we observed significant enhancer activity (FDR <0.01) for 118 tiles comprising 71 distinct peaks. The greatest activity was observed in the promoter region and in the first intronic enhancer identified in our previous study^7^. Weaker activity was observed throughout the tested interval. Only 12 tiles comprising 10 distinct peaks demonstrated significant repressor activity. Coordinates in peaks from the CRISPRn and CRISPRi screens were modestly more likely than those in non-peaks to overlap an MPRA active tile (odds ratios = 2.4 and 2.6, respectively), but the same was not true for the CRISPRa screen (odds ratio = 1.0, Fig 5B).

### Integrated analysis of CRE screening modalities

Comparing the results of our CRISPR nuclease, inhibition, and activation screens and MPRA, we next examined the overlap in the regions identified by each approach (Figs 6, 7A). Only the promoter region was consistently identified by each approach. As discussed above, the CRISPRn and CRISPRi screens exhibited a high degree of concordance, with coordinates in peaks from each also showing a modest ∼2-3-fold greater odds than those in non-peaks of overlapping an active tile by MPRA. CRISPRa peaks, by contrast, were largely unique. Only those in the promoter colocalized with CRISPRn or CRISPRi peaks, and coordinates in CRISPRa peaks were no more likely than those in CRISPRa non-peaks to overlap an MPRA active tile.

**Figure 6.**
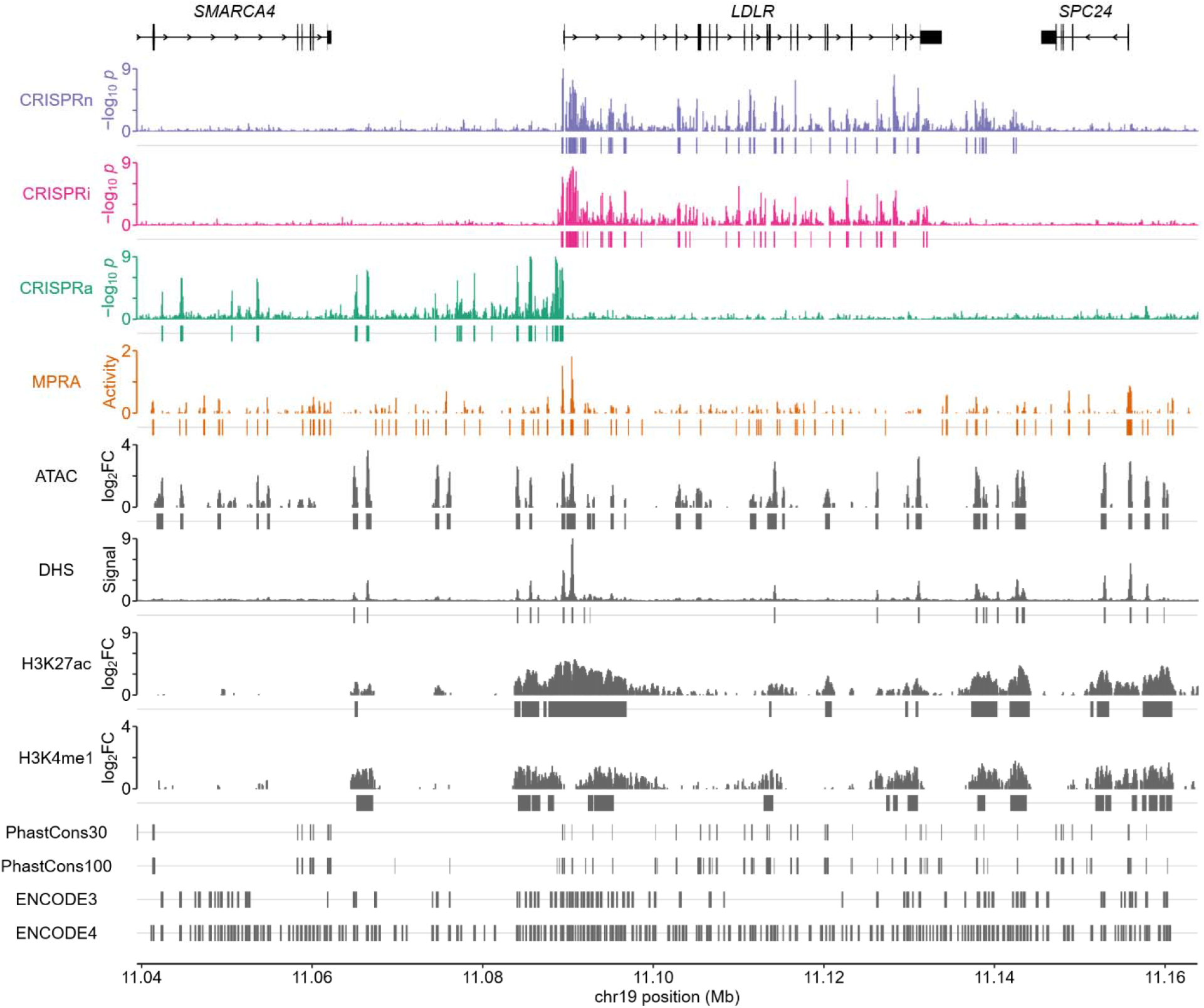
Spatial relationship of *LDLR* screen data to predictors of CRE activity. Genomic tracks for enhancer activity and peak calls as assessed by each screening modality (LDL^high^ population gRNA depletion in the CRISPRn and CRISPRi screens, gRNA enrichment in the CRISPRa screen, and increased activity in the MPRA) visualized in comparison to integrated signals for open chromatin (ATAC-seq and DHS-seq) and histone marks (H3K27ac and H3K4me1 ChIP-seq) as well as regions identified with evolutionary conservation (PhastConsElements 30-way primate and 100-way vertebrate tracks) and candidate CRE activity (ENCODE3 and ENCODE4 releases).

**Figure 7.**
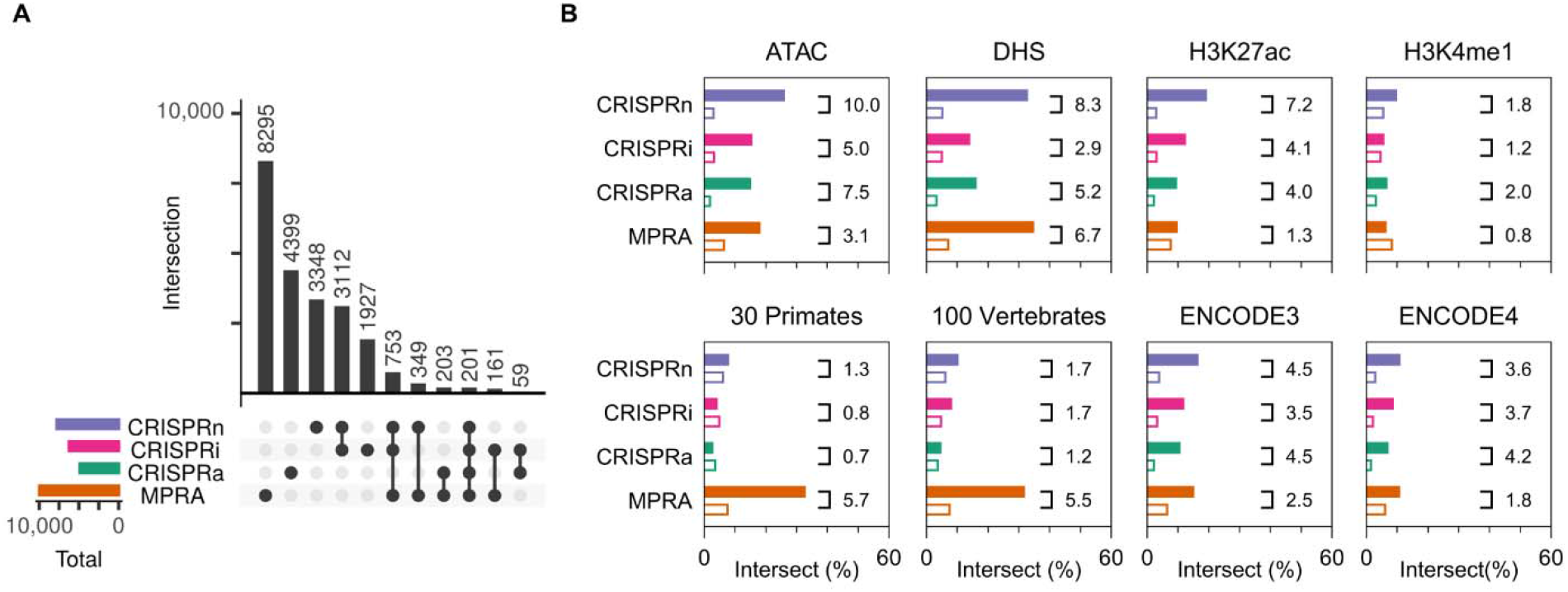
Predictive performance of enhancer-associated features for functional activity. (A) Upset plot of the number of coordinates identified with functionally significant enhancer activity by each screening modality. (B) For each indicated classifier, coordinates were stratified based on whether they were predicted to have CRE activity or not. The proportion of coordinates in each group overlapping a functionally significant enhancer region, as determined by each screening modality, is displayed along with their corresponding odds ratio.

We also assessed the performance of different CRE-associated features in predicting functional activity by each screening modality (Fig 7B, Supplementary Table 6). The strongest correlation was observed for ATAC-seq, whose peaks were enriched in hits from all screening modalities but tracked most closely with CRISPRn (odds ratio = 10.0). DHS-seq likewise exhibited the strongest correlation with CRISPRn (odds ratio = 8.3). More modest predictive performance was apparent for H3K27ac and H3K4me1 ChIP-seq, PhastCons evolutionary conservation scores, and ENCODE cCRE calls. For each predictor of CRE activity, the majority of identified regions were not supported by empirical evidence of functional significance in any assay. An interactive web viewer with the screening data and associated genomic features in Figure 6 is available at https://www.emmerlab.org/LDLR-noncoding.

## DISCUSSION

Pioneering studies of *LDLR* gene regulation used episomal reporters to map sterol response elements (SREs) in the promoter that mediate the sensitivity of *LDLR* expression to cellular sterol concentrations^11-13^. Subsequent work identified the transcription factors (SRE-binding proteins, or SREBPs) mediating their activity^14-17^ and their binding sites across the genome^18-20^. Other investigations have implicated binding sites in the *LDLR* promoter for additional transcription factors including Sp1^21-23^, CREB^24^, and C/EBP-β^24,25^. More recent systems genetics approaches have further identified MAFF and ATF3 as regulators of *LDLR* expression with ChIP-based evidence of promoter binding^26,27^.

Outside of the promoter, only a few putative *LDLR* enhancers have been identified with evidence of functional activity. In the 3’ intergenic space, the “PS2” enhancer was hypothesized due to its evolutionary conservation and validated by reporter assays^28^. In the first intron, one study found an impact of specific human LDL-associated variants on reporter activity, but did not detect enhancer activity for the major allele^29^; our prior study observed robust activity of the major allele and confirmed relevance at the endogenous locus via enhancer deletion^7^. Several other *LDLR* noncoding regions have also been identified with features suggestive of CRE activity but have not been empirically tested.

In this context, we now provide a comprehensive map of perturbation effects and reporter activity across the *LDLR* locus. Each of our approaches demonstrated reproducibility, readily detected control sequences, and identified new regions with strong statistical signals. Notably, although some of these regions had been predicted to harbor activity based on biochemical or evolutionary features, several were not. Likewise, multiple regions with predicted CRE activity demonstrated no significant effects in our functional assays, consistent with other studies finding imprecision in CRE predictions^6^.

Our identification of functionally significant *LDLR* CREs has several potential applications. Loss of function *LDLR* mutations are the most common cause of Familial Hypercholesterolemia (FH), an autosomal dominant Mendelian disease affecting an estimated 1 in 250 people worldwide and characterized by early onset ASCVD. The pathogenic mutation spectrum of *LDLR* is dominated by coding variants^30^, with a few exceptions in the proximal promoter and 5’ UTR^31^ or deep intronic splice regions^32,33^. However, the relative lack of CRE-localized pathogenic variants may reflect an ascertainment bias, as distal regulatory regions are rarely interrogated in standard FH genetic screening. We anticipate that the regions implicated by our study may help prioritize noncoding variants for future investigations to determine whether they either cause FH or incrementally affect ASCVD risk, depending on the magnitude of their effect. Moreover, identifying functionally significant variants that increase rather than disrupt *LDLR* expression may nominate candidates for somatic genome editing, an area of active development for LDL-targeted therapeutics^34^.

Aside from the implications of our findings for *LDLR* specifically, our study also serves as a prototype for the application of CRISPRn, CRISPRi, CRISPRa, and MPRA interrogation to the same noncoding space. Several notable patterns emerged. The regions identified by CRISPRn and CRISPRi screens were highly concordant. We initially suspected that the distribution of CRISPRn peaks in primarily intronic regions might be due to a subset of edited cells with larger indels affecting nearby coding sequences and/or splice sites. Indeed, a prior study using paired gRNA deletions identified such a mechanism driving enrichment signals^35^. With the important caveat that we filtered CRISPRn target sites within 100 bp of intron-exon junctions, we found that most CRISPRn peaks were replicated by CRISPRi screening with a nuclease-deficient Cas9 that silences CREs without indel formation. While disruption of splicing might still occur by nuclease-independent steric hindrance, the lack of similar effects in CRISPRa screening argues against this mechanism and suggests that our CRISPRn and CRISPRi peaks arose due to genuine disruption of enhancer activity. Interestingly, although most intronic peaks identified by CRISPRn were also called by CRISPRi, a subset of functional elements in the 3’ intergenic space including the established PS2 enhancer^28^ were only detected by CRISPRn. This pattern of substantial but incomplete overlap is broadly consistent with prior studies of *BCL11A*^36^ and a panel of genes in the DNA mismatch repair pathway^37^.

In contrast to the general concordance of our CRISPRn and CRISPRi peaks, the regions identified by our CRISPRa screen were almost entirely distinct. The basis for this discordance is unclear but likely reflects the underlying mechanism of each modality, with CRISPRn and CRISPRi identifying elements that are necessary for basal expression and CRISPRa identifying elements that are not individually necessary for basal expression but sufficient to augment expression when activated. Unique CRISPRa peaks may therefore represent functionally redundant CREs, such that their disruption is compensated for while their forced activation still increases expression. Alternatively, an element may be latent or poised under basal culture conditions, with no native activity to disrupt but retained capacity to respond to recruited activators, as described for stimulus-responsive immune enhancers^38^. The lack of enrichment in CRISPRa peaks for MPRA active tiles may support this model of latent or poised activity. We also cannot exclude the possibility that CRISPRa peaks represent synthetic interactions between the promoter and regions without true endogenous activity. Conversely, CRISPRn/CRISPRi peaks may go undetected by CRISPRa if dCas9-VP64 displaces native transcription factors without exceeding their activating function^39^. At enhancers, this competition for occupancy has been associated with weaker activation than native transcription factor binding alone^40^. Regardless of the underlying mechanisms, significant discordance between CRISPRa and CRISPRn/CRISPRi does not appear unique to *LDLR*, as it has been similarly reported for other CREs^41,42^.

Finally, our MPRA tiling of the *LDLR* locus provided complementary information on which individual sequences were able to drive expression outside their native genomic context. MPRA active tiles were modestly enriched for CRISPRn and CRISPRi peaks but not for CRISPRa peaks, suggesting that elements contributing to basal expression are more likely to retain autonomous, sequence-intrinsic activity than the elements identified only by forced activation. MPRA active tiles were also enriched for ATAC-seq and DHS-seq signal and for evolutionary conservation, consistent with a link between native chromatin accessibility, sequence constraint, and sequence-intrinsic regulatory capacity. However, we found the majority of MPRA active tiles to localize to regions with no significance by any of the CRISPR screening modalities. As MPRA is increasingly used as a tool to nominate candidate regulatory variants, our findings suggest that further perturbation testing at the endogenous locus may help refine which MPRA-active variants are most likely to have a truly causal effect on gene expression.

Several limitations to our approach warrant consideration. All CRISPR screens are inherently sensitive to the availability of PAM sites at a target region and the variable efficiencies of gRNAs targeting them. Our MPRA library consisted of staggered and overlapping 129 bp tiles, which may not detect effects for larger CREs that are not amenable to a reductionist approach. Likewise, our MPRA vector uses a synthetic minimal promoter that may not capture CRE effects that are specific to the *LDLR* promoter. Our functional assays relied on HuH7 cells. Although these are a widely used model for hepatocytes that recapitulate major pathways of *LDLR* regulation^43,44^, it is likely that individual *LDLR* CREs will have cell type-specific effects.

These caveats notwithstanding, our study represents the most comprehensive functional map of the noncoding *LDLR* space to date, systematically identifying discrete regulatory regions and highlighting the strengths and limitations of each approach to their identification. The regions we define focus the mechanistic and variant-level studies needed to establish their individual contributions to *LDLR* expression and ASCVD risk.

## Acknowledgments

This research was supported by the National Institutes of Health R01-HL167733 (BTE), R01-HL171013 (BTE), the A. Alfred Taubman Medical Research Institute (BTE), the American Heart Association 24POST1198445 (HSV), and the Michigan Pioneer Fellows Program (HSV). The funders had no role in study design, data collection and analysis, decision to publish, or preparation of the manuscript.

## Author Contributions

HSV and BTE conceived the project. All authors contributed to experimental design and execution, data analysis, and manuscript preparation.

## MATERIALS AND METHODS

### Reagents

HuH7 cells (JCRB Cell Bank, Osaka Japan) were cultured in DMEM (Thermo Fisher Scientific, Waltham MA) supplemented with 10% fetal bovine serum (Sigma-Aldrich, St. Louis MO) and penicillin-streptomycin (Thermo Fisher Scientific). Cells were maintained in a humidified 37°C incubator with 5% CO2. Oligonucleotide sequences (Integrated DNA Technologies, Coralville IA) are provided in Supplementary Table 7. Individual CRISPR gRNA constructs were prepared by T7 ligase insertion of phosphorylated and annealed oligonucleotides into BsmBI-digested (New England Biolabs, Ipswich MA) lentiGuide-Puro^45^ (Addgene, Watertown MA, #52963) or pXPR_502^8^ (Addgene, #96923). Additional plasmids were obtained for expression of SpCas9^45^ (Addgene, #52962), dCas9-KRAB^46^ (Addgene, #96918), and dCas9-VP64^47^ (Addgene, #61425). Plasmids were verified by whole plasmid sequencing (Plasmidsaurus, South San Francisco CA) and lentiviral stocks were prepared in coordination with the University of Michigan Vector Core.

### CRISPR library design and synthesis

Preparation of a custom gRNA library for pooled lentiviral expression was adapted from prior studies^7,43,48,49^. In brief, every SpCas9 gRNA sequence within the ∼124 kb region of interest (GRCh38/hg38 chr19:11,039,463–11,163,820) was identified based on its NGG protospacer adjacent motif using the CRISPR Targets track of the UCSC Genome Browser Table Browser^50,51^. Control gRNAs comprised 100 non-targeting sequences, repurposed from a prior study^52^, and 20 gRNAs apiece for *LDLR* and *MYLIP*, each selected using CRISPick with 10 apiece optimized for CRISPR interference or CRISPR activation^8,53^. Every gRNA sequence was appended with invariant flanks for primer annealing and synthesized as a pooled oligonucleotide library (GenScript, Piscataway NJ). PCR amplification was performed using Herculase II (Agilent Technologies, Santa Clara CA) and primers that provided vector homology arms, with cycle number optimized based on amplicon analysis with a TapeStation D1000 Assay (Agilent Technologies). HiFi assembly (New England Biolabs) was performed using 125 ng of amplicon and 825 ng of BsmBI-digested lentiGuide-Puro. Assembly products were purified using a QIAquick PCR purification kit (QIAGEN, Hilden Germany) and electroporated into NEB 10-beta electrocompetent *E. coli* (New England Biolabs) using a MicroPulser (Bio-Rad Laboratories, Hercules CA) Ec1 setting. Serial dilutions of electroporated bacteria were spread onto 10 cm LB agar plates to quantify total colony counts and compared to control assemblies of digested vector only to confirm >20:1 colony ratios. Plasmid DNA was extracted from all colonies scraped from 245 mm LB agar plates using a Plasmid Plus Midi kit (QIAGEN). PCR amplification of plasmid pools was performed with the primers designated in Supplementary Table 7 that provided flanking barcodes and sequencing adapters. Massively parallel sequencing was performed in the University of Michigan Advanced Genomics Core with an Illumina MiSeq Micro paired-end 2 x 150 bp run, with individual gRNA sequences mapped and enumerated using MAGeCK^54^. Analysis of mapped reads confirmed >99.9% coverage with a ratio of <10 for the 90^th^ and 10^th^ percentile most abundant gRNAs.

### Cellular LDL uptake

HuH7 cells were first transduced with lentiviral constructs expressing Cas9, dCas9-KRAB, or dCas9-VP64, selected 1 day later with 4 µg/mL blasticidin (Thermo Fisher), and cultured for 14 days. A second transduction was then performed with either individual gRNA constructs for pilot studies or with the pooled gRNA library at MOI ∼0.5 and >500X library coverage for high throughput screening. A second round of selection was then performed 1 day later with 3 µg/mL puromycin (Thermo Fisher) and cells were propagated for an additional 14 days. Transduced cells were incubated with 4 µg/mL DyLight549-conjugated LDL (Cayman Chemical, Ann Arbor MI) in serum-free DMEM for 2 hours, detached with TrypLE Express (Thermo Fisher), and washed and resuspended in phosphate-buffered saline with 2% FBS. Flow cytometry was performed on a Ze5 Analyzer (Bio-Rad Laboratories) for single guide testing and on a FACS Discover S8 (BD, Franklin Lakes NJ) for screen bulk sorting of a minimum of 500,000 cells apiece representing 10% subpopulations with the most and least fluorescent LDL uptake. All CRISPR screens were performed for 3-4 independent biologic replicates. Genomic DNA was purified from sorted cells using a DNeasy kit (QIAGEN). Quantification of gRNA abundance in each sample was performed by massively parallel sequencing of PCR amplicons, as described above, with an Illumina NovaSeq paired-end 2 x 150 read. Individual reads were mapped to gRNA sequences using MAGeCK^54^ and are provided in Supplementary Tables 2-4.

### CRISPR screen analysis

CRISPR screens were analyzed using SPANA (Saturating Perturbation Analysis of Noncoding Activity; v1.0.0; see Code availability). Differential gRNA abundance was analyzed using DESeq2^55^. Guides not reaching at least 10 counts in at least 3 samples were removed. Preliminary analysis using non-targeting guides as the null set revealed a subtle but reproducible offset from targeting guides, consistent with nonspecific effects of double-strand break formation rather than diffuse regulatory activity across the locus^56,57^. We also attempted normalization across the entire test guide pool, but under default median-of-ratios normalization most guides were displaced from zero rather than centered on it, suggesting compositional asymmetry in a library where genuinely depleted guides outnumber enriched ones. We therefore estimated size factors from a data-adaptive null set identified by iterative trimming of the LFC distribution (SPANA-null), following the trimming logic of TMM normalization^58^ but with a convergence-based threshold and retaining median-of-ratios estimation for consistency with the downstream dispersion model. Dispersions were estimated once and held fixed across passes. Within each pass, guides more than 2 SD from the mean LFC were iteratively excluded until membership stabilized. The resulting null set was supplied as controlGenes to re-estimate size factors and passes continued until the Jaccard similarity of null set membership between consecutive passes reached at least 0.98, which occurred within three passes in all screens. Control guides were excluded from the test population and analyzed separately under the converged size factors. A final Wald test was run under the converged size factors without Cook’s distance outlier filtering. Both maximum-likelihood and apeglm-shrunken LFC estimates^59^ were calculated. One-sided p-values for depletion and enrichment were derived from the normal cumulative distribution of the Wald statistic and each corrected for multiple hypothesis testing using the Benjamini-Hochberg method^60^. Spatial clustering of significant guides was assessed by permutation (SPANA-cluster), comparing the median nearest-neighbor distance among guides passing a Benjamini-Hochberg adjusted p-value of 0.01 to a null generated over 100,000 permutations by reassigning hit labels among all tested guide positions. One-sided empirical p-values were computed as the proportion of permutations at or below the observed value. Per-guide apeglm-shrunken LFC estimates were averaged across sliding 100 bp genomic windows (SPANA-scan), with one window centered on each test guide. Because guide density varies across the locus, empirical significance was assessed against a size-matched null. For each observed window size, 10^9^ window means were simulated by drawing the same number of scores with replacement from the test guide score distribution, and one-sided empirical p-values for enrichment and depletion were computed as the proportion of simulated means at or beyond the observed mean, floored at the simulation resolution of 10^-9^. Overlapping windows with a Benjamini-Hochberg adjusted empirical p-value at or below 0.01 were merged into contiguous peaks.

### MPRA library design and synthesis

The general strategy for lentiMPRA synthesis was adapted from prior studies^7,61^. For the same region of interest interrogated by the saturating CRISPR screens (GRCh38/hg38 chr19:11,039,463–11,163,820), a total of 2,899 overlapping 129 bp tiles, staggered every 43 bp, were derived from the hg38 reference sequence downloaded from UCSC Genome Browser^50^. Control tiles were selected to include 100 random sequences as well as sequences previously tested in HepG2 cells^61^, including the 50 synthetic sequences and 50 natural enhancers with the greatest activity and the 50 synthetic sequences with the least activity. All control sequences were included in both forward and reverse orientations. *LDLR* test tiles and control tiles were appended with invariant flanks, synthesized as a pooled oligonucleotide library, PCR amplified with primers listed in Supplementary Table Ft7 that also provided vector homology arms and a random 15 bp 5’ UTR barcode, and analyzed by TapeStation D1000 assay (Agilent Technologies). HiFi assembly (New England Biolabs) was performed with 150 ng of amplicon and 850 ng of AgeI/SbfI-digested (New England Biolabs) pLS-SceI^61^ (Addgene, # 137725). Assembly products were column purified, electroporated, and resulting plasmids harvested as described above for CRISPR library synthesis. PCR amplicons of tile sequences and their corresponding 5’ UTR barcodes were generated using the primers listed in Supplementary Table 7. Tile representation and barcode associations were analyzed with an Illumina NextSeq paired-end 2 x 150 run with custom sequencing primers. Titers of resulting lentiviral stocks were calculated by qPCR from cells transduced with a range of virus volumes, as previously described^61^.

### MPRA testing and analysis

For each of 8 independent biologic replicates, ∼8 million HuH7 cells were seeded in pairs of 15 cm plates at approximately ∼25% confluence and transduced with the pooled lentiviral library at MOI ∼10. Cells were harvested 2 days later and split into equal parts for isolation of genomic DNA and RNA using DNeasy and RNeasy Plus Mini kits, respectively (QIAGEN). To remove residual genomic DNA, 60-120 µg of each RNA sample was treated with 12 units of TURBO DNAse (Thermo Fisher) for 30 ming at 37°C. Conversion of RNA to cDNA was performed using the MPRA-RT-P1 primer and the SuperScript III First-Strand synthesis kit (Thermo Fisher) per manufacturer’s instructions. Genomic DNA and cDNA samples were used as templates for PCR amplification of the CRS-barcode sequences with the primers listed in Supplementary Table 7. The abundance of the CRS barcode in each sample was quantified by sequencing on an Illumina NovaSeq paired-end 2x150 bp run. Barcode counts were extracted from FASTQ files and aggregated to their corresponding tiles based on the tile-barcode association map described above. Activity for each tile was quantified using MPRAnalyze^10^ with depth factors estimated per replicate, then standardized against the median and median absolute deviation of random control sequences and assessed by two-sided testing with Benjamini-Hochberg correction. Tiles with an adjusted p-value below 0.01 were called significant, and those with overlapping or abutting genomic intervals were merged into contiguous peaks.

### Comparative analysis

CRISPR screen and MPRA peaks corresponding to enhancer activity (gRNA depletion in CRISPRn and CRISPRi screens, gRNA enrichment in CRISPRa screens, and increased tile activity in MPRA) were compared to publicly available data for chromatin accessibility (human liver ATAC-seq and DHS-seq) and histone modifications (human liver H3K27ac and H3K4me1 ChIP-seq)^62^. To visualize signal across multiple independent experiments of the same type, log2-transformed values at each coordinate were extracted from bigWig files (accessions in Supplementary Table 6) using WiggleTools^63^ (v1.2.11) with an offset of 0.01 and averaged. Consensus peaks were defined as coordinates falling within called peaks in a minimum of 50% of contributing experiments. Evolutionarily conserved elements (PhastCons 30-way primate and 100-way vertebrate^64^) and candidate CREs nominated by ENCODE3^65^ and ENCODE4^5^ were obtained from the UCSC Table Browser^51^. For each comparison data set, base pairs across the region of interest (excluding those within 100 bp of a coding sequence) were classified as falling within or outside its annotated elements, and the likelihood of overlapping a CRISPR screen or MPRA peak was compared between the two groups as an odds ratio.

### Data availability

Guide RNA library annotations, mapped read counts, and analysis results for each CRISPR screen are provided in Supplementary Tables 1–4, and MPRA data in Supplementary Table 5. Accession numbers for the public ENCODE datasets used are listed in Supplementary Table 6. An interactive browser of these data is available at https://emmerlab.org/LDLR-noncoding.

### Code availability

The SPANA code used in this study (version 1.0.0), comprising SPANA-null, SPANA-scan, and SPANA-cluster, is freely available at https://github.com/emmerlab/spana and archived on Zenodo at https://doi.org/10.5281/zenodo.23545216. The code is written in R and released under the GNU General Public License, version 2 or later. Source code for the interactive browser is available at https://github.com/emmerlab/LDLR-noncoding-browser under the GNU General Public License, version 3 or later.

---

**Figure S1.**
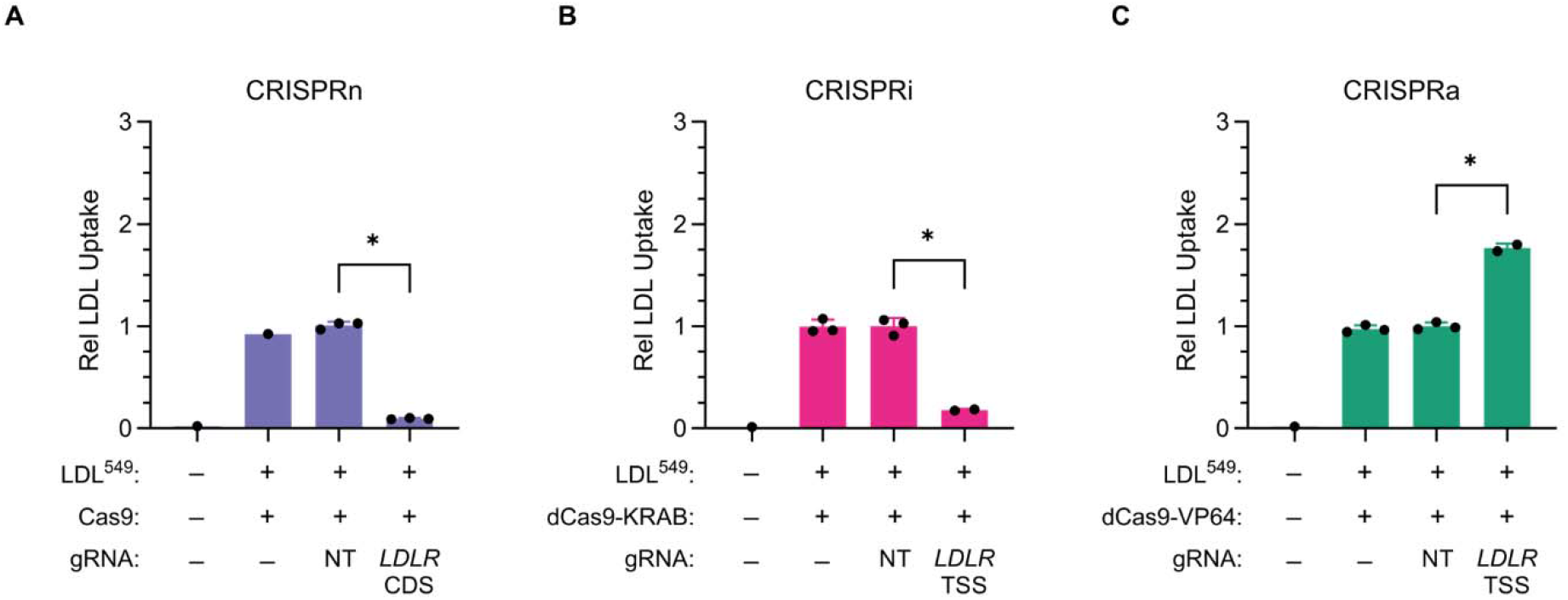
Modulation of cellular LDL uptake by different mechanisms of CRISPR-based *LDLR* perturbation. Relative mean fluorescence intensity (MFI) corresponding to uptake of exogenous fluorescent LDL by HuH7 cells transduced with Cas9 nuclease (A), dCas9-KRAB (B), or dCas9-VP64 (C) in combination with the indicated gRNAs.

**Figure S2.**
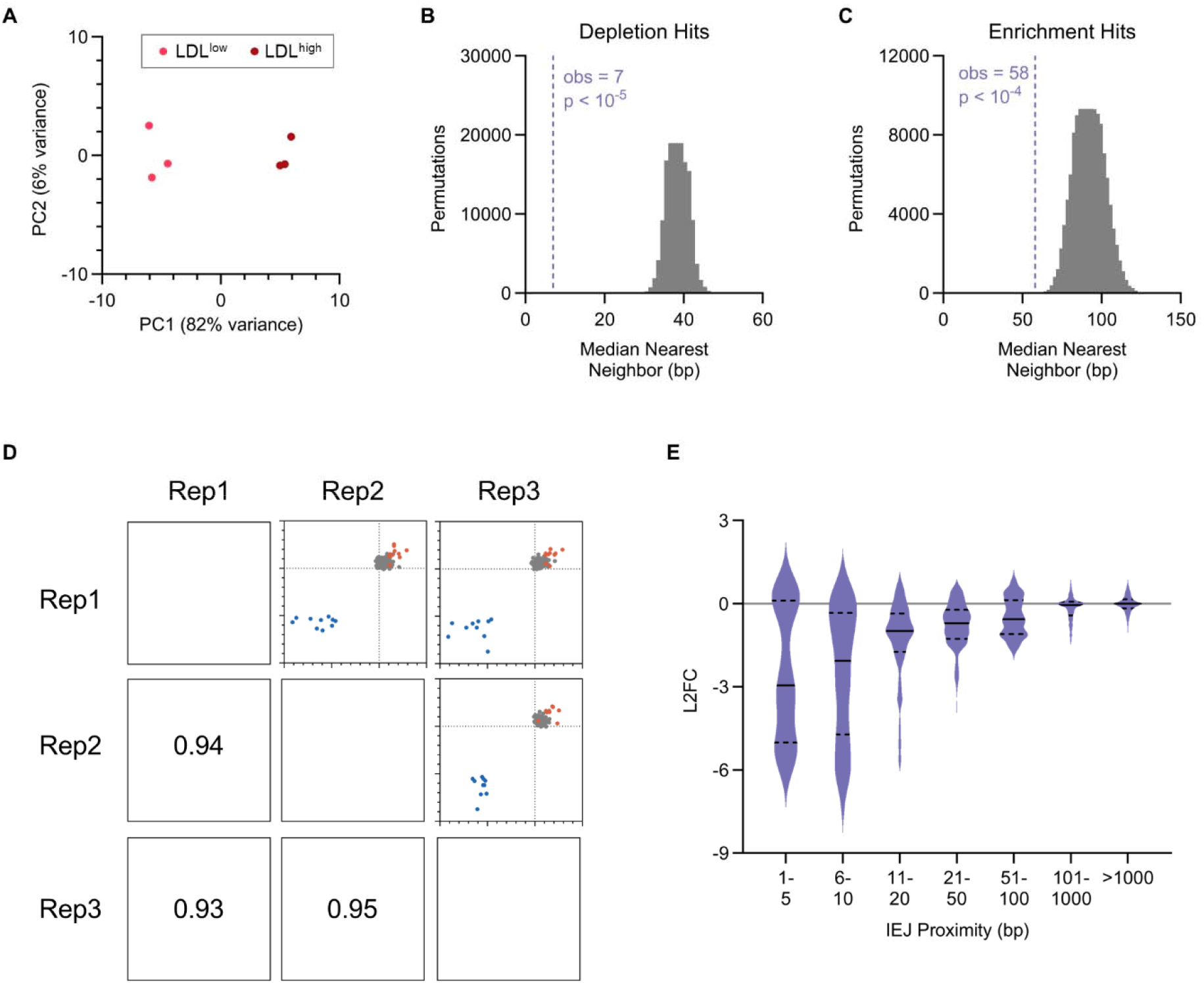
Quality control analysis of CRISPR nuclease screen. (A) Principal component analysis of gRNA counts in sorted LDL^high^ and LDL^low^ subpopulations over 3 independent biological replicates. (B) Nearest neighbor testing for significantly depleted gRNAs, with the vertical line representing the observed median proximity from one hit to the next closest hit and the distribution representing the same proximity after random shuffling of gRNA effects over 100,000 permutations. (C) Nearest neighbor testing for significantly enriched gRNAs. (D) Pairwise correlation of gRNA log2FC of control gRNAs over independent biologic replicates. Values represent Pearson correlation coefficients. (E) Violin plot of distribution of gRNA log2FC for gRNAs grouped by their proximity to the nearest intron-exon junction (IEJ). Solid lines indicate medians and dashed lines indicate upper and lower quartiles.

**Figure S3.**
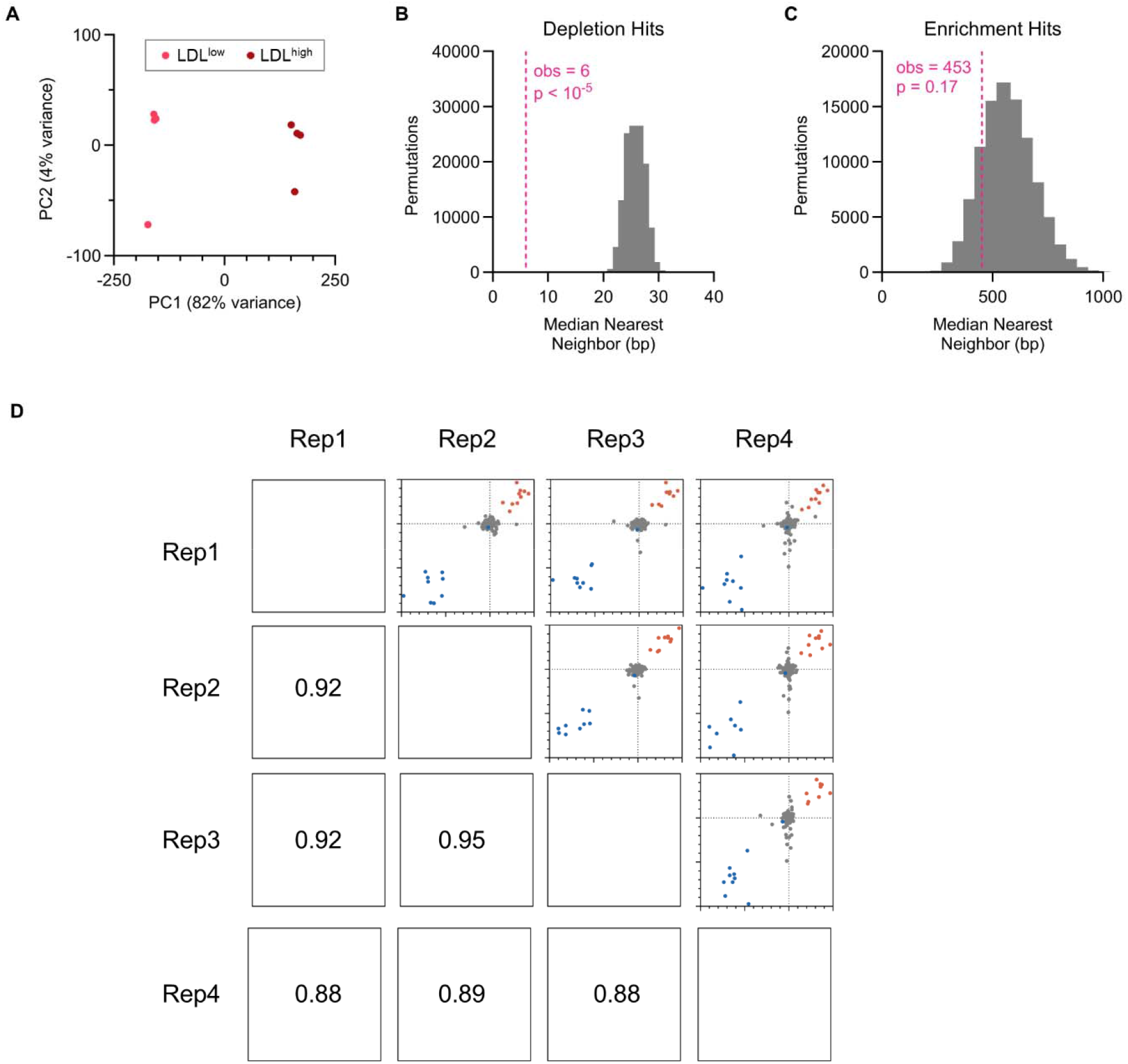
Quality control analysis of CRISPR inhibition screen. (A) Principal component analysis of gRNA counts in sorted LDL^high^ and LDL^low^ subpopulations over 4 independent biological replicates. (B) Nearest neighbor testing for significantly depleted gRNAs, with the vertical line representing the observed median proximity from one hit to the next closest hit and the distribution representing the same proximity after random shuffling of gRNA effects over 100,000 permutations. (C) Nearest neighbor testing for significantly enriched gRNAs. (D) Pairwise correlation of gRNA log2FC of control gRNAs over independent biologic replicates. Values represent Pearson correlation coefficients.

**Figure S4.**
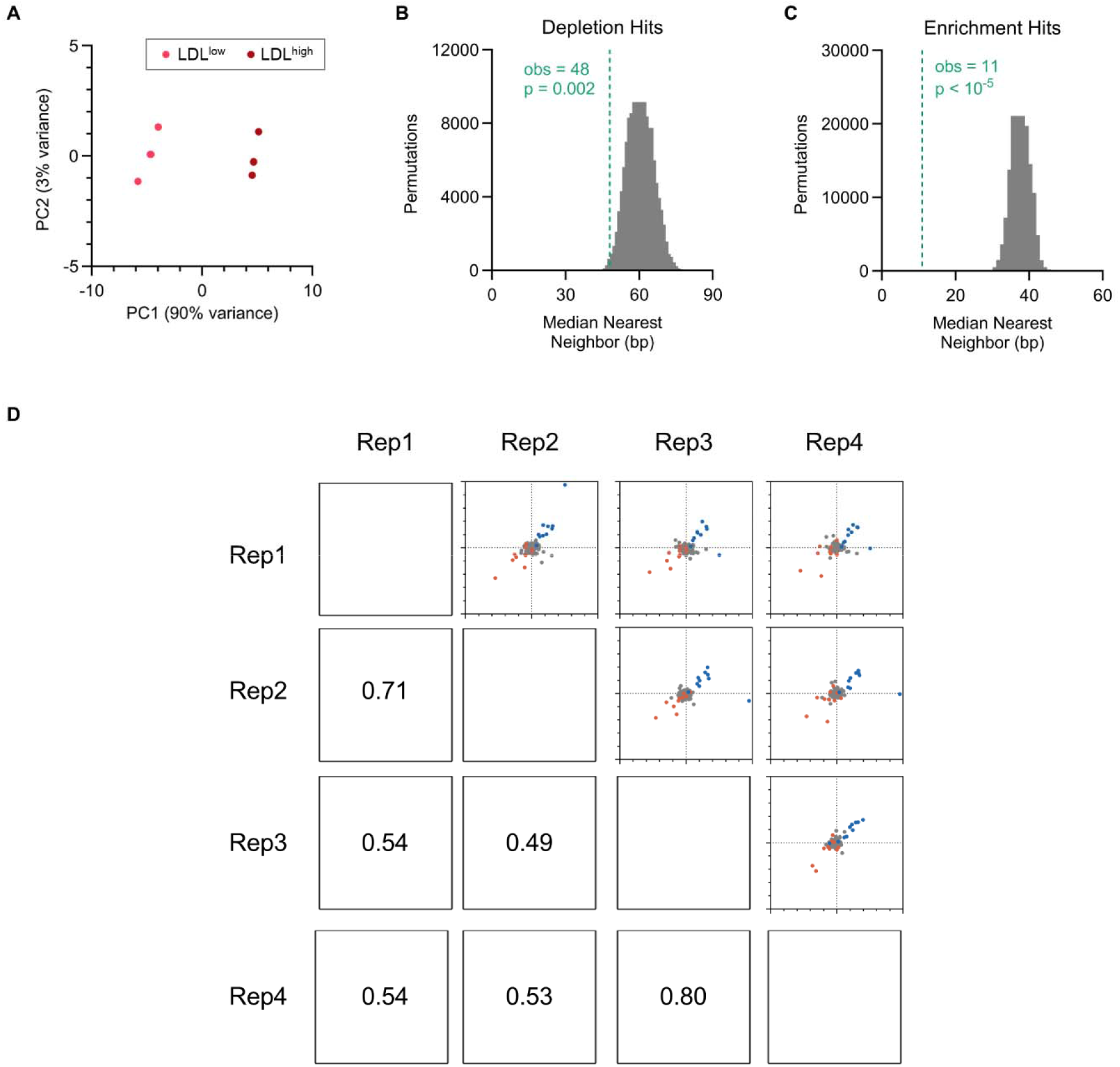
Quality control analysis of CRISPR activation screen. (A) Principal component analysis of gRNA counts in sorted LDL^high^ and LDL^low^ subpopulations over 4 independent biological replicates. (B) Nearest neighbor testing for significantly depleted gRNAs, with the vertical line representing the observed median proximity from one hit to the next closest hit and the distribution representing the same proximity after random shuffling of gRNA effects over 100,000 permutations. (C) Nearest neighbor testing for significantly enriched gRNAs. (D) Pairwise correlation of gRNA log2FC of control gRNAs over independent biologic replicates. Values represent Pearson correlation coefficients.

**Figure S5.**
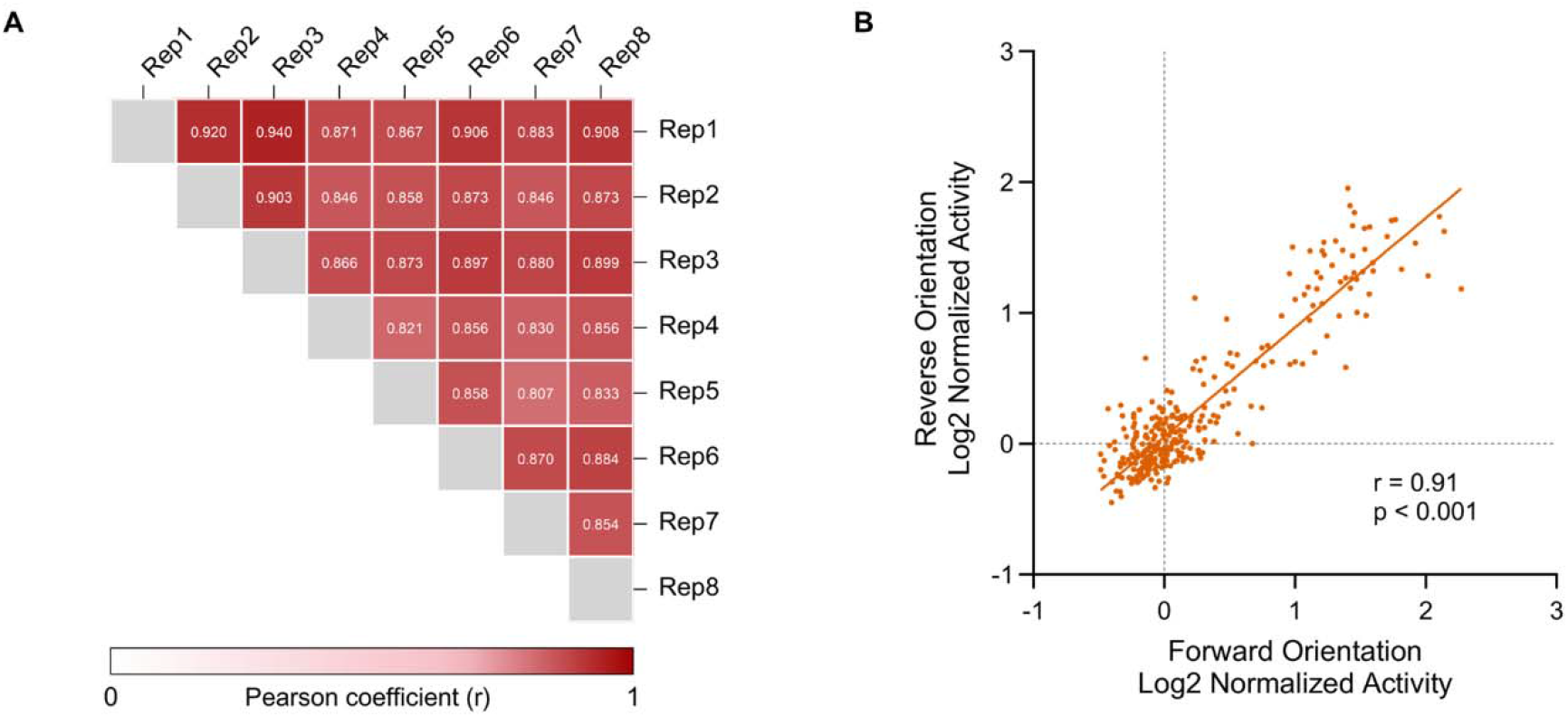
Quality control analysis of MPRA screen. (A) Matrix of Pearson coefficients for control tile activity between pairwise combinations of independent biologic replicates for the MPRA study. (B) Correlation between normalized activity of control tiles assayed in forward and reverse orientation.

**Supplementary Table 1. CRISPR library composition.** A unique identifier is provided for each guide sequence, derived from its estimated DSB coordinate and strand when used for nuclease screening. Guides of type “test” target noncoding regions more than 100 bp from the nearest *LDLR* CDS and are used to identify the null set for DESeq2 normalization.

**Supplementary Table 2. CRISPR nuclease screen data.** Raw counts, individual gRNA enrichment analysis, sliding 100 bp window enrichment analysis, and peak calls for gRNA abundance in HuH7 LDL^high^ relative to LDL^low^ sorted cells in the pooled CRISPR nuclease screen.

**Supplementary Table 3. CRISPR inhibition screen data.** Raw counts, individual gRNA enrichment analysis, sliding 100 bp window enrichment analysis, and peak calls for gRNA abundance in HuH7 LDL^high^ relative to LDL^low^ sorted cells in the pooled CRISPR inhibition screen.

**Supplementary Table 4. CRISPR activation screen data.** Raw counts, individual gRNA enrichment analysis, sliding 100 bp window enrichment analysis, and peak calls for gRNA abundance in HuH7 LDL^high^ relative to LDL^low^ sorted cells in the pooled CRISPR activation screen.

**Supplementary Table 5. Massively parallel reporter assay data.** Aggregate raw barcode counts in HuH7 genomic DNA and mRNA and derived activity scores and peak calls for each tile in the MPRA library.

**Supplementary Table 6. Analysis of CRE-predictive features.** ENCODE accession information, mean signal, and consensus peaks calls for publicly available human liver ATAC-seq, DHS-seq, and H3K27ac and H3K4me1 ChIP-seq experiments.

**Supplementary Table 7. Oligonucleotide sequences.**

